# Identification of Subsets of Genetic Alterations in KRAS-mutant Lung Cancer using Association Rule Mining

**DOI:** 10.1101/235390

**Authors:** Junior Tayou

## Abstract

**Background:** Lung cancer is the leading cause of all cancer death accounting for 1 out of 4 cancer-related death in both men and women. KRAS mutations occur in ~ 25% of patients with lung cancer, and the presence of these mutations is associated with poor prognosis. Efforts to directly target KRAS or associated downstream MAPK or the PI3K/AKT/mTOR pathways have seen little or no benefits. One probable reason for the lack of progress in targeting KRAS-mutant tumors is the co-occurrence of other cell survival pathways and mechanisms.

**Method and results:** To identify other potential cell survival pathways in subsets of KRAS-mutant tumors, I performed unsupervised machine learning on somatic mutations in metastatic lung cancer from 725 patient samples. I identified 67 other genes that were mutated in at least 10% of the samples with KRAS alterations. This gene list was enriched with genes involved in the MAPK, AKT and STAT3 pathways, cell-cell adhesion, DNA repair, chromatin remodeling, and the Wnt/beta-catenin pathway. I also identified 160 overlapping subsets of 3 or more genes that code for oncogenic or oncosuppressive proteins that were mutated in at least 10% of KRAS-mutant tumors.

**Conclusions:** In this study, I identified genes that are co-mutated in KRAS-mutant lung cancer. I also identify subpopulations of KRAS-mutant lung cancer based on the set of genes that were also altered in the tumor samples. The design of research models that captures these subsets of KRAS-mutant tumors would enhance our understanding of the disease and facilitate personalized treatment for lung cancer patients with KRAS alterations.

## Background

Advanced lung cancer is the most commonly diagnosed, and the leading cause of all cancer death worldwide [1, 2]. Morphologically, lung cancer can be broken down into two major categories: non-small cell lung carcinoma (NSCLC) which makes up 85% of all the cases and small cell lung carcinoma (SCLC) [2]. Data accumulated over the last 15 years shows that lung cancer is not a single disease but a collection of genetically defined neoplasms with distinct molecular features and clinical outcomes. The known oncogenic drivers of lung cancer include the receptor tyrosine kinases (RTKs): EGFR, ALK, RET, MET, DDR2, ROS1 and FGFR1 and their downstream facilitators KRAS, PIK3CA, and BRAF [3]. Tumors with EGFR alterations are targetable with EGFR tyrosine kinase inhibitors (TKIs) including erlotinib, gefitinib, and afatinib [4–6]. Rearrangement of ALK and ROS1 genes are targeted with crizotinib and ceritinib[7]. Progress has also been made in targeting mutations in the other RTKs [3, 8]. However, for tumors containing alterations in KRAS, there are no FDA approved drugs. KRAS mutations are present in between 20 and 25% of lung cancer and are associated with poor prognosis [9, 10]. KRAS signal primarily through the MAPK and the PI3K/AKT/mTOR pathways [11–13]. Efforts to directly target KRAS or its downstream effectors have seen little or no benefit [14]. The reason for the lack of success in targeting KRAS effectors might be due to the development of adaptive response mechanisms or the co-occurrence of mutations in other genes that activate different cell survival pathways in KRAS-mutant tumors.

Here, I employed unsupervised machine learning using association rules to identify genes whose mutations co-occurs with KRAS alterations in advanced lung cancer. The building blocks of association rules are the items that may appear in any given transaction [15]. This approach is largely applied in supermarkets to learn about consumer purchasing patterns. Retailers use it to identify products that are usually purchased together in other to tailor their advertisement outreach. Transactions are specified in terms of itemsets. For example, in a grocery store, {Bread, peanut butter, jelly} represent a typical itemset. A rule from this transaction might be expressed in the form {peanut butter, jelly} → {bread}, in other words, if peanut butter and jelly (left-hand side or LHS) are purchased together, then bread (right-hand side or RHS) is also likely to be bought. Supermarkets apply association rules to find “interesting” combinations of items that are not as apparent as the peanut butter, jelly and bread example.

I used the Apriori association rule algorithm [16], which states that all subsets of a frequent itemset must also be frequent. For example, for the rule {mutation X} → {mutation Y}, then the items {mutation X}, {mutation Y} and {mutation X and mutation Y} must all be frequent. The strength of a rule is determined by three statistical measures: the support, confidence and lift statistics. The support of a mutation or mutation set is a measure of how frequently it occurs in the tumor samples. The support for one or more mutations in a gene X in N tumor samples is given by

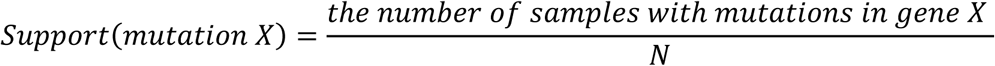

The confidence is the measure of the accuracy of a rule. For a given rule {mutation X} → {mutation Y}, the confidence is defined as

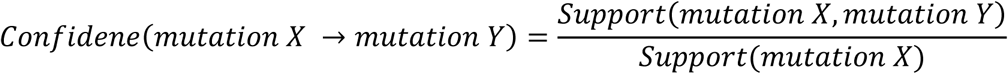

The confidence tells us the proportion of the samples where a mutation in gene X results in a mutation in gene Y. The lift statistic measures how much more likely a mutation or a set of mutations is present in a sample relative to its typical rate of occurrence, given that another mutation or set of mutations are present in the sample. The lift for a given rule {mutation X} → {mutation Y} is define by

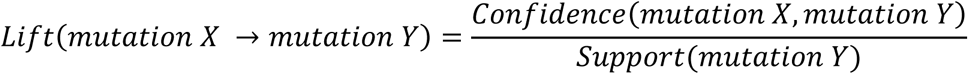

For a hypothetical example, {TP53, BRACA1} → {CD43}, support = 0.2, confidence = 0.80, lift = 3, count = 125. This rule can be interpreted as – if a tumor has mutations in both TP53 and BRCA1, it will also contain a mutation in CD43. This rule covers 20% (support = 0.2) of all the tumor samples, and the rule is correct in 80% (confidence = 0.80) of tumors having TP53 and BRCA1 mutations. A lift of 3 implies that samples with mutations in both TP53 and BRCA1 are three times more likely to have a mutation in CD43 than a typical cancer sample.

I applied the Apriori machine learning algorithm to a dataset containing somatic mutations from 725 advanced lung cancer patient samples to identify the subsets of genes that were also altered in tumors with KRAS mutations.

## Methods

### Data acquisition

The data used in this study was obtained from the CBioportal website. The data represents the findings from a study on the mutational landscape of advanced cancer from more than 10,000 patients by Zehir A et al. [17]. I downloaded the data on October 6th, 2017. After extraction of all the information on mutations including deletion, insertion, substitution and gene rearrangements from all metastatic lung cancer samples, I merged the columns of interest by sample ID. The resulting file containing the genes with alterations was converted into a sparse matrix (supplementary table 1). I converted the genes coding for receptor tyrosine kinases (PTPRT, PTPRD, and PTPRS) to PTPRs and the B melanoma antigen precursors (BAGE 2, 3, 4, and 5) to BAGE.

### Apriori algorithm

Association rule was performed using the Apriori algorithm that was developed by Agrawal R and Srikant R [16]. I used the “arules” and “aruleviz” packages in the R programming language to execute the apriori algorithm and for visualization of the resulting rules respectively. For a typical analysis, I used the codes:

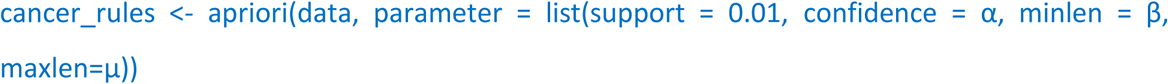

I used a support of 0.01 in all the analyses; this threshold will include only the genes that were mutated in at least 1% of the samples. More than 600 genes were above this threshold. The values for the confidence, the minimum (minlen) and the maximum (maxlen) number of the members in a rule were selected based on the question that was addressed.

### Graph analysis

Rules or subrules generated using the Apriori algorithm in R were saved as Gephi file and manipulated in Gephi 0.9.2 to generate directed graphs as shown in figures 2D and 4A. For example, the rule

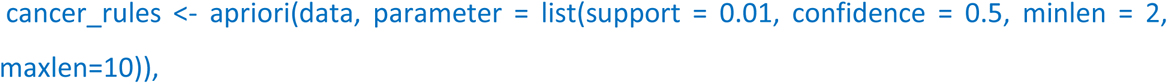

can be save as a Gephi file using the code:

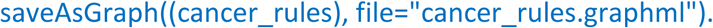

In Gephi, graphs were manipulated to represent nodes with different colors or labels with different font sizes based on the directionality and magnitude of the degree of connectivity of the nodes.

### Bioinformatics

I used the REACTOME pathway analysis site to determine the pathways that were enriched in the gene sets obtained from association rule analysis.

## Results and discussion

### Distribution of mutations of known oncogenic drivers in metastatic lung cancer

The dataset used in this study contained 725 metastatic lung cancer patient samples (93% NSCLC and 7% SCLC)[17]. Consistent with previous studies [18], TP53 was mutated in 63% of all metastatic lung cancer samples. ROS1 and ALK1 mutations were present in 17% and 15% of the tumors respectively. EGFR alterations were present in 29% of the samples, while DDR2, RET, MET and FGFR1 were mutated in 8%, 12%, 9% and 6% respectively. The RTK facilitators PIK3CA, BRAF and KRAS, were altered in 8%, 16% and 23% of the samples respectively. The samples that had KRAS mutations were almost exclusively NSCLC; only 0.6% (1 tumor sample) was SCLC. All the tumors that had EGFR mutations were NSCLC.

### Analyses of rules statistics

Using a support of 1% and confidence of 10%, I identified 187,168 rules for the genes that were mutated in all the samples. KRAS mutations were associated with 5,724 of these rules. The supports for the 187,168 rules was in the range of 1 and 25% with most of the rules covering between 1 and 10% of the data (Figure 1A). Rules containing KRAS had a narrower support range. The supports for the KRAS-mutant subrules were predominantly between 1 and 5%. The maximum support for a rule that had a KRAS alteration was 12% (Figure 1B). The confidence for all the rules and the KRAS-mutant subrules were evenly spread between 10 and 100% (Figure 1A, B).

**Figure 1:**
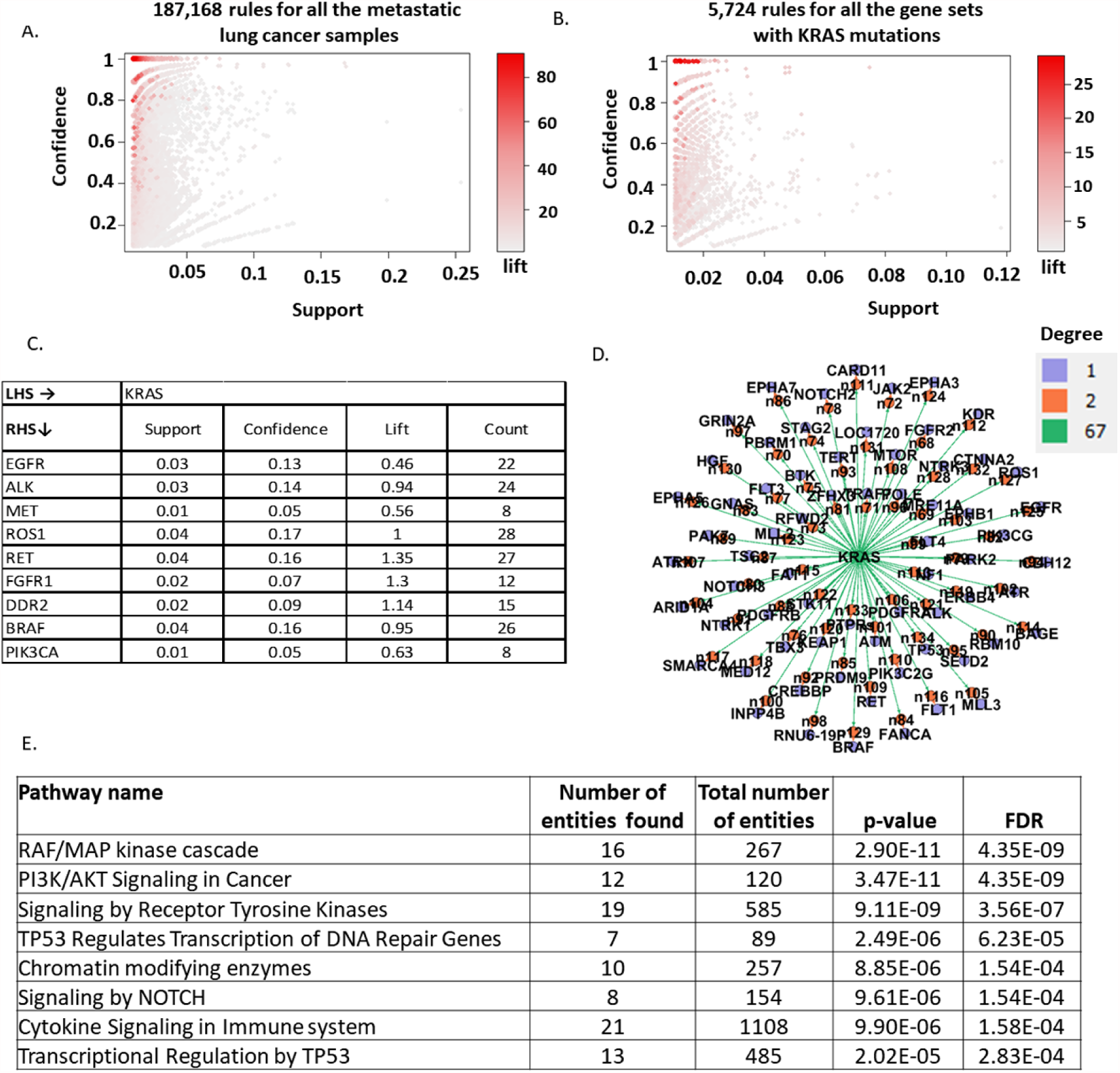
KRAS-mutant lung cancer is associated with alterations in genes that regulate diverse cellular processes: Scatter plot showing the distribution of (A) all the rules generated for the genes that were mutated in all the metastatic lung cancer samples. (B) The subset of the rules containing KRAS. (C) The association between mutations in KRAS and other tumor drivers. Note that the lift (likelihood) for KRAS-mutant tumors to also have mutations in EGFR, MET or PI3K3CA was less than one would expect by chance. (D). A directed graph showing the association between KRAS and 67 genes. I limited our query to a one to one association and with KRAS on the LHS to identify the genes that were also mutated in KRAS-mutant samples. I used a confidence of 0.1 to include only the genes that were altered in at least 10% of KRAS samples. The subrules generated in R was imported into Gephi and manipulated to represent the nodes with distinct colors based on their degree of connectivity. (E) I used the 67 genes that were changing as input into the REACTOME pathway analysis tool. Shown in E, is a subset of the pathways that were enriched in the 67-member gene list.

### KRAS-mutant cancers were less likely to have EGFR, MET or PIK3CA mutations

I examined the co-occurrence of mutations in KRAS and alteration in other oncogenic drivers in metastatic lung cancer. Consistent with previous reports [19], EGFR and KRAS mutations were almost mutually exclusive; only 3% (support = 0.03) of the tumors had mutations in both oncogenes (Figure 1C). The likelihood for a sample with a KRAS alteration to also have an EGFR mutation was two-fold (lift = 0.46) less than one would expect in a typical metastatic lung cancer sample (Figure 1C). Similarly, the likelihood for mutations in MET (found in 1% of KRAS-mutant samples) or PIK3CA (1% of KRAS-mutant samples) to occur in KRAS-mutant samples were less than one would expect by chance. The likelihoods that a sample with alterations in KRAS will also have mutations in ALK, ROS1, RET, FGFR1, DDR2 or BRAF were all close to 1 (Figure 1C).

### Mutations in KRAS were associated with alterations in 67 other genes

To identify potential cell survival pathways that might also be activated in KRAS-mutant tumors. I queried the rules that I generated from the Apriori algorithm for genes associated with KRAS mutations. I identified 67 genes that were mutated in at least 10% of the KRAS-mutant tumors (Figure 1D and Table 1). These genes are involved in diverse biological pathways that are important for cancer cell survival (Figure 1E). Genes involved in RTK signaling and the downstream MAPK, PI3K/AKT/mTOR and the JAK/STAT pathways were predominantly altered (Figure 1E, 2A). This list was also enriched with genes coding for proteins involved in chromatin remodeling and transcriptional regulation by TP53 (Figure 1E, 2C).

**Figure 2:**
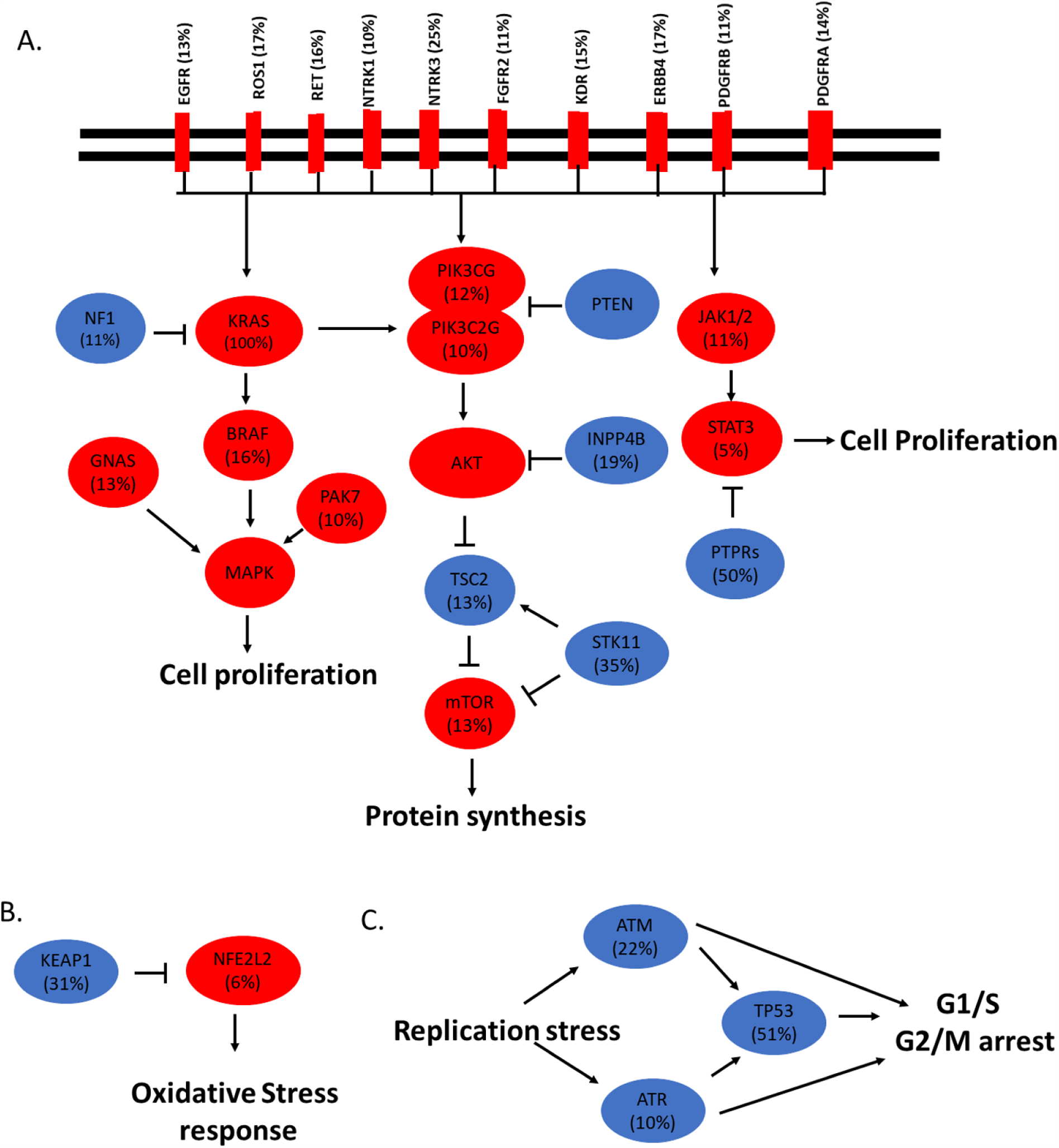
Schematics of genes in the MAPK, mTOR and STAT3 pathways, the oxidative stress response and the DNA repair mechanisms that were altered in KRAS-mutant lung cancer: Schematics showing the relationship between the genes that were mutated in KRAS-mutant tumors. Red denotes oncogenic proteins, and blue is for oncosuppressive proteins. (A) The members of the MAPK, PI3K/AKT/mTOR, and STAT3 pathways, (B) KEAP1 regulation of oxidative stress response, (C) TP53/ATM/ATR mediated cell cycle arrests.

**Table 1:**
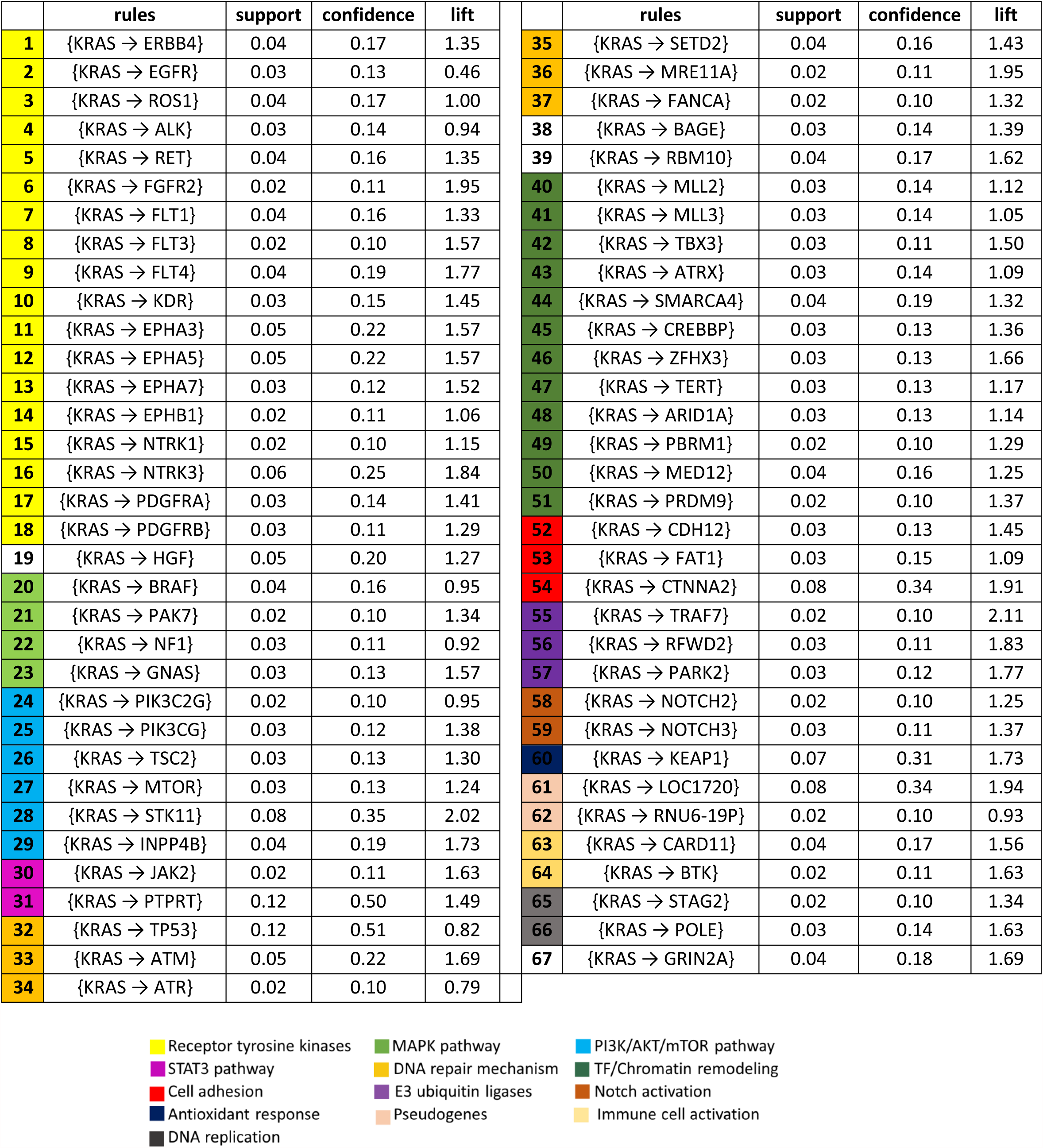
Genes associated with KRAS in metastatic lung cancer: The support is the proportion of metastatic lung cancer covered by the rule. Confidence (accuracy) is the proportion of the KRAS-mutant tumors covered by the rule. Lift is the likelihood of having a mutation in the gene on the right of the rule given that the sample has an alteration in KRAS.

### Mutation of RTK genes in samples with KRAS mutations

In addition to the known oncogenic RTK drivers discussed earlier, I identified several other RTKs that were altered in KRAS-mutant tumors including members of the EGFR family, the EPH, NTRK, and VEGF receptor families. ERBB4, a member of the EGFR family, was mutated in 17% of the samples with KRAS alterations (Figure 2A). Unlike EGFR (Figure 1C), the likelihood of having a tumor with mutations in ERBB4 given that the tumor has an alteration in KRAS was higher than one would expect by chance in an advanced lung cancer sample (Table 1, lift = 1.35). KRAS-mutant samples also had alterations in several members of the VEGF receptor family including FLT1 (16%), FLT3 (10%), FLT4 (19%) and KDR (15%) receptors (Figure 2A, Table1). The NTRK1 and NTRK3 that codes for TrkA and TrkC receptors were mutated in 10% and 25% of KRAS-mutant samples respectively (Figure 2A, Table 1). Four members of ephrin receptor family: EPHA3 (22%), EPHA5 (22%), EPHA7 (12%) and EPHB1 (11%) were also altered in KRAS-mutant tumors (Table 1). The platelet-derived growth factor receptors, PDGFRA and PDGFRB, were altered in 14% and 11% of the KRAS-mutant samples respectively (Figure 2A, Table 1). Mutations in MET and KRAS were almost mutually exclusive (Figure 1C). However, MET’s ligand HGF was mutated in 20% of KRAS-mutant samples (Table 1).

### Mutation of genes involved in the MAPK and PI3K/AKT/mTOR pathways

KRAS is an effector of RTK signaling and mutation in KRAS can lead to constitutive activation of the downstream cell survival MAPK pathway irrespective of receptor activation [20, 21] (Figure 2A). Consequently, RTK inhibitors have been shown to be futile in the presence of activating KRAS mutations [20, 21]. BRAF is a serine-threonine kinase directly downstream of KRAS in the MAPK pathway (Figure 2A). Constitutively active BRAF mutations can also drive tumor progression by activating MEK irrespective of the status of RTK and KRAS [22]. In colorectal cancer, it has been shown that mutations in KRAS and BRAF are almost mutually exclusive [23, 24]. However, in the metastatic lung cancer dataset used in this study, BRAF was mutated in 16% of samples with KRAS mutations (Figure 1C). Also, the likelihood of having a KRAS-mutant tumor with a BRAF alteration was 0.95 (Table 1). Activating mutations in the Gαs subunit of heterotrimeric G-proteins can also promote cancer cell proliferation via the MAPK pathway [25]. I observed that GNAS which codes for the Gαs subunit was mutated in 13% of the KRAS-mutant samples (Figure 2A). The tumor suppressor NF1 was altered in 11% of KRAS-mutant tumors (Figure 2A). NF1 activates the GTPase activity of KRAS [26]. However, loss of NF1 is not redundant in KRAS-induced myeloproliferative disorders as alterations in both genes resulted in a disease with a shorter latency [27]. These observations suggest that NF1 might have additional oncosuppressive functions and the presence of a concurrent mutation in KRAS might lead to a more aggressive disease. In summary, I identified four genes downstream of receptor activation with genetic alterations in the MAPK pathway.

RTKs and KRAS signaling can also activate the PI3K/AKT/mTOR pathway [13]. This pathway is commonly activated and genetically altered in lung cancer [28, 29]. I observed that alterations of several components of the PI3K/AKT/mTOR pathway were present in KRAS-mutant tumors. PIK3CG, PIK3C2G, TSC2, and mTOR were mutated in 12%, 10%, 13% and 13% of the tumors with KRAS alterations respectively (Figure 2A and Table 1). Consistent with previous studies [29], I observe that mutations of individual components of this pathway were almost mutually exclusive (Data not shown). The tumor suppressor, INPP4B was mutated in 19% of KRAS-mutant tumors (Figure 2A and Table 1). PI(3,4)P2 is essential for AKT activation. INPP4B inactivates AKT by dephosphorylating PI(3,4)P2 [30]. Also, alterations in INPP4B can drive tumorigenesis via activation of serum and glucocorticoid-regulated kinase 3 (SGK3) [31]. STK11 is a serine/threonine kinase that functions as a tumor suppressor by regulating TP53-dependent apoptosis pathways [32] and also by activating TSC1/2 in the PI3K/AKT/mTOR pathway [33, 34] (Figure 3A). STK11 is commonly mutated in lung cancer [19], and it was mutated in 17% of all metastatic lung cancer samples in our dataset (Data not shown). Consistent with prior studies [19, 35], I observed that STK11 mutations were two times more likely to occur in KRAS-mutant cancer as opposed to KRAS wild-type tumors (Table 1). STK11 alterations were found in 35% of tumors with KRAS-mutant tumors. On the contrary, STK11 mutations were three times less likely to occur in EGFR-mutant tumors as opposed to EGFR wild-type cancer samples. Only 6% of the EGFR-mutant tumor had alterations in STK11 (Data not shown). These observations are consistent with the fact that STK11 and KRAS mutation occurs predominantly in smokers while EGFR mutations occur mostly in tumors from patients without a smoking history [19]. STK11 also seems to have additional effects in KRAS-mutant tumors. Mutations in STK11 was shown to suppress the immune surveillance in KRAS-mutant lung cancer, and they might serve as a predictor of de novo resistance in immune checkpoint blockade [36, 37]. In summary, I identified at least six genes downstream of RTK activation that were mutated in the PI3K/AKT/mTOR pathway.

**Figure 3:**
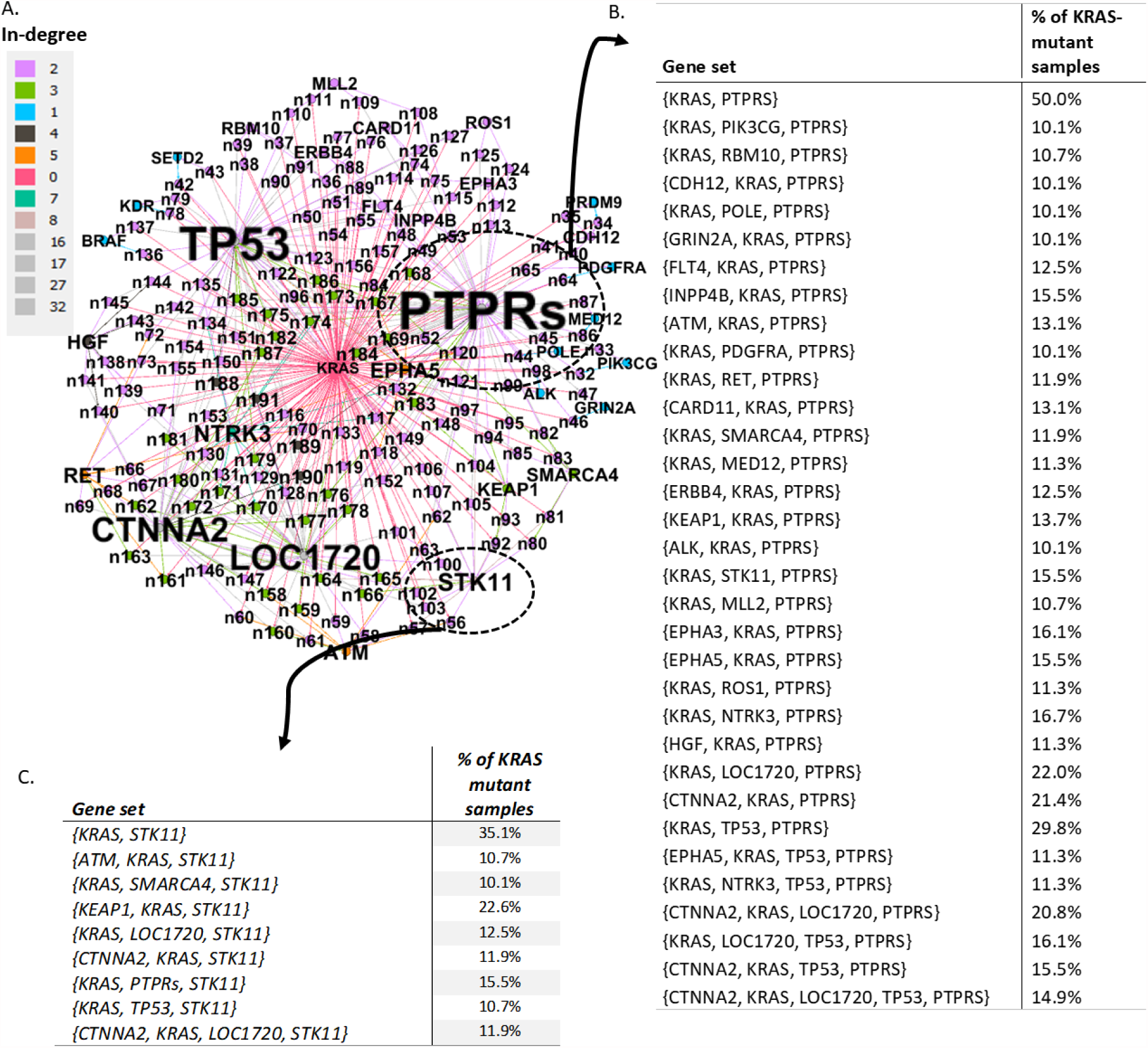
Analyses of rules with 3 or more members: (A) I generated all the rules that contained KRAS as in figure 1B. The rules were then subsetted to include only rules with at least three genes, with KRAS on the LHS and that was present in at least 10% of KRAS-mutant samples. The resulting 160 rules were imported as a graph file into Gephi. Files were manipulated to show the notes of different colors based on the edges coming into the nodes (in-degree). Note that KRAS had an indegree of zero since it was restricted to the LHS. Mutations in PTPRs were the most connected with 32 in-degrees. (B) and (C) shows the tabular representation of the rules that converge on PTPRs and STK11 nodes.

### Mutations in protein receptor tyrosine phosphatases and STAT3 pathway

The receptor-type protein tyrosine phosphatases (PTPRT, PTPRD, and PTPRS) were mutated in more than 50% of KRAS-mutant samples. These tumor suppressors were two times more likely to occur in samples with KRAS mutations as opposed to tumors with wild-type KRAS (Table 1). On the other hand, PTPRs mutations were less likely to be present in samples with EGFR alteration (data not shown). Consistent with previous studies [38, 39], PTPRT was mutated in 17% of all metastatic lung cancer samples, while PTPRD and PTPRS were mutated in 18% and 7% of the samples respectively. At least one of these PTPRs was mutated in 33% of all samples (Table 1). The role of PTPRs in lung cancer is yet to be unraveled. However, phospho-STAT3 is a substrate for these phosphatases [40, 41] and frequent loss of function mutations in PTPRT and PTPRD have been linked to hyperphosphorylation of STAT3 in head and neck squamous carcinoma and in gliomagenesis [42–44]. STAT3 is hyperactive in some lung cancers [45]. This hyperactivity might be due to enhanced signaling from RTKs, cytokines or JAK kinases (altered in 11% of KRAS-mutant tumors) or by inactivating mutations in PTPRs [42, 46, 47].

### Alterations in the antioxidant stress response pathway

Another tumor suppressor gene that had a high rate of genetic alterations in KRAS-mutant samples was KEAP1. Concurrent mutations in KRAS and KEAP1 in lung cancer is associated with a more aggressive disease [48]. The KEAP1 gene encodes for Kelch-like ECH associated protein 1 which is a negative modulator of NFE2L2, a master transcriptional regulator of the antioxidant response. Loss of KEAP1 was shown to alter the response of KRAS-mutant lung cancer cells to MAPK inhibitors. Krall E et al. showed that loss of KEAP1 could abolish the increase in reactive oxygen species that are associated with MAPK inhibition. They went further to show that deletion of KEAP1 can change the cell metabolism and enhance the survival and proliferation of the lung cancer cells in the absence of MAPK signaling [49]. In another study by Romero R et al., lung cancer cells with concurrent mutations in KRAS and KEAP1 were shown to be dependent on increase glutaminolysis and were more sensitive to glutaminase inhibition [50]. I observed that KEAP1 was 1.7 times more likely to be mutated in KRAS-mutant as oppose to KRAS-wildtype tumors. I observed that 30% of KRAS-mutant samples had mutations in KEAP1 (Figure 2B and Table 1). On the contrary, EGFR-mutant tumors were more than two-fold less likely to have a mutation in KEAP1 (data not shown).

### Mutations in DNA replication stress response genes

Replication stress is an attribute of many malignant cells [51]. ATR mediate response to stress by arresting the progression of the cell cycle while stabilizing and repairing the replication fork. ATM and TP53 maintain the integrity of the genome when ATR is defective. ATM and TP53 arrest cell cycle progression allowing for DNA repair [52, 53]. I observed that TP53, ATM, and ATR were altered in 51%, 22% and 10% of the samples with KRAS mutations respectively (Table 1 and Figure 2C). ATM and ATR mutations were almost mutually exclusive in KRAS-mutant cells, and none of the samples with concurrent mutations in {KRAS, ATM, ATR, TP53}. However, {ATM, KRAS, TP53} and {ATR, KRAS, TP53} subsets were present in 9.5% and 6% of KRAS-mutant samples (Data not shown). KRAS-mutant samples also had alterations in SETD2 (16%), FANCA (10%) and MRE11 (11%) which codes for proteins that regulate repair of DNA mismatch and double-strand breaks. In summary, six genes that are involved in DNA repair and the replication stress response mechanisms were mutated in at least 10% KRAS-mutant samples.

### Mutations in transcription factors and chromatin remodeling genes

KRAS-mutant tumors had alterations SMARCA (19%), ARIDIA (13%), PBRM1 (10%), PRDM9 (10%), CREBBP (13%), TBX3 (11%), ATRX (14%), MLL2 (14%), MLL3 (14%), TERT (13%), ZFHX3 (13%) and MED12 (16%) genes that codes for transcription factors and proteins involved in chromatin remodeling (Table 1). MED12 promotes tumorigenesis in part by regulating the Wnt/beta-catenin pathway [54]. I also identified mutations in other genes that are important for the formation of the beta-catenin/TCF transactivating complex including MLL2, MLL3, CREBBP, TERT, and SMARCA. SMARCA, ARIDIA, and PBRM1 are part of the chromatin remodeling complex SWI/SNF. The genes that encode for the subunits of the SWI/SNF complex are altered in more than 20% of all human cancers. Deficiency in one or more of these subunits including ARIDIA is associated with enhanced tumor growth and invasiveness [55]. Taken together, 12 genes coding for transcription factors and chromatin modifying proteins that were altered in at least 10% KRAS-mutant tumors.

### Mutations in E3 ubiquitin ligases

Another group of genes that were altered in KRAS-mutant samples were genes coding for E3 ubiquitin ligases. PARK2, TRAF7, and RFWD2 were mutated in 12%, 11% and 10% of the samples with KRAS alterations (Table 1). E3 ubiquitin ligases mediate the degradation of many proteins that are important for cancer development [56]. PARK2 deletion or loss of function was shown to promote inflammation, genomic instability, and progression of lung cancer [57, 58]. TRAF and RFWD2 can ubiquitinate and promote degradation of TP53 leading to decrease TP53-dependent transcription and apoptosis [59]. In KRAS-mutant samples, mutations of the individual E3 ubiquitin ligases were mutually exclusive.

### Mutations in Cadherins and β-catenins

Cadherins and catenins control cell-cell adhesion and dysregulation of their expressions are associated with increased metastasis and poor prognosis of lung and other cancers [60, 61]. I observed that CDH12 and the atypical cadherin FAT1 were genetically altered in 13% and 15% of KRAS-mutant tumors respectively (Table 1). Mutations in FAT1 and CDH12 were mutually exclusive in KRAS-mutant tumors. CTNNA2 which codes for α2-catenin was mutated in 34% of KRAS-mutant samples (Table 1). KRAS-mutant tumors were twofold more likely to have a mutation in CTNNA2 than KRAS wild-type tumors. Mutations in CTNNA2 were less likely to occur in EGFR samples; only 10% of EGFR samples had alterations in CTNNA2, and half of these samples had concurrent {CTNNA2, EGFR, KRAS} mutations (Data not shown). The effect of CTNNA2 dysfunction have not been addressed in lung cancer, but frequent mutations in CTNNA2 were shown to promote migration and invasiveness of head and neck squamous cell carcinoma cells [62]. The mutation subsets {KRAS,CTNNA2,FAT1} and {KRAS,CTNNA2,CDH12} were present in 5% and 8% of KRAS-mutant tumors respectively. In summary, cadherins and catenins were collectively altered in at least 40% of KRAS-mutant tumors.

### Analyses of rules involving three or more genes

To understand the complexity of the sub-populations of KRAS-mutant tumors, I queried the rules that were generated for the subtypes of cancers with mutations in three or more genes. As shown in figure 1B, mutations in KRAS were associated with 5724 rules. When I limited the query to rules with at least three genes and that were present in at least 10% of KRAS-mutant tumors, 160 subrules were generated (Figure 3A and Table S2). Most (79%) of these rules had mutations in three genes. Many of these rules were associated with mutations in PTPRs, TP53, CTNNA2 and the pseudogene LOC1750 (Figure 3A). There was a great degree of “overlap” between the rules. For example, all the 17 rules with CTNNA2 on the RHS were similar to the 95% of the rules with LOC1750 on the RHS.

Mutations in PTPRs can lead to constitutive activation of STAT3 which is associated with many lung cancer [42–44]. PTPRs alterations were collectively altered in at least 50% of KRAS-mutant tumors (Table 1). A concurrent mutation in KRAS might lead to a tumor with hyperactive MAPK, AKT and STAT3 signaling [20, 21]. I explored the subsets of tumors with KRAS and PTPRs mutations to identify other potential pathways that might be defective (Figure 3B). I identified {KRAS, PTPRT, SMARCA4}, {KRAS, PTPRT, MED12}, and {KRAS, PTPRT, MLL2} subsets in 11.9%, 11.3% and 10.7% of KRAS-mutant tumors respectively. In addition to hyperactivity of MAPK, AKT and STAT3 pathways, additional mutations in SMARCA4, MED12 or MLL2 could also implicate Wnt/beta-catenin pathway and transcriptional regulation by the SWI/SNF complex [54, 55]. Another subset of tumors with KRAS and PTPRT mutations were {KRAS, KEAP1, PTPRT} and {CTNNA2, KRAS, PTPRT} in 14% and 21.4% of the KRAS-mutant tumors respectively. Concurrent mutations in KRAS, KEAP1, and PTPRT can lead to a tumor with hyperactive MAPK, AKT and STAT3 pathways and defective antioxidant response mechanisms [49, 50]. While mutations in CTNNA2, KRAS, and PTPRT could lead to a tumor with defective cell-cell adherence in addition to constitutive activation in the MAPK, AKT and STAT3 pathways [62].

STK11 alterations are associated with poor prognosis in lung cancer [35]. However, there are conflicting reports on the effects of alterations in STK11 in KRAS-mutant tumors. Some researchers reported that concurrent STK11 and KRAS mutations did not affect the aggressiveness of the disease in patients [48, 63]. Findings from other studies show that STK11 can potentiate tumor progression in KRAS-mutant tumors in mouse models [64] and was associated with a more aggressive disease [36]. It is possible that this discrepancy might be due to the presence of other alterations in subpopulations of KRAS^mut^/STK11^mut^ lung cancer. I observed that about 66% of KRAS^mut^/STK11^mut^ tumors also had alterations in KEAP1, which regulates oxidative stress response [49, 50]. I also observed that STK11^mut^/PTPRs^mut^ and STK11^mut^/CTNNA2^mut^ were present in about 35%, and 33% of the tumors with KRAS mutations (Figure 3C). Such combinations of mutations might affect the biology of the disease leading to different clinical outcomes. The design of research models that can capture such combinations of mutation could enhance our understanding of the disease.

### Conclusion

In this study, I have implemented a smart approach to finding subpopulations of metastatic KRAS-mutant lung cancers based on the combination of somatic mutations in genes coding for oncogenic and oncosuppressive proteins. Designing research models that captures these overlapping subpopulations could lead to the development of combination therapies that are effective and can circumvent potential resistance mechanisms in KRAS-mutant metastatic lung cancer.

## Declarations

### Ethics approval and consent of participants

Not applicable

### Consent for publication

Not applicable

### Availability of data and materials

The data supporting the findings from in this study were obtained from the CBioportal site[17]. I have included supplementary files for all the analyses that I performed.

## Acknowledgement

Not applicable

## R codes

#Load data and required packages

library(arules)

library(arulesViz)

data <-read.transactions("data.csv", sep = ",")

#Model training

\## Support = 0.01, confidence = 0.1 and minimum length of rule = 2

cancer_rules <-apriori(data, parameter = list(support = 0.01, confidence = 0.1, minlen = 2))

#Figure 1A

plot(cancer_rules)

\# Figure 1B

KRAS_rules <-subset(cancer_rules, subset = items %ain% c(“KRAS”))
plot(KRAS_rules)

\# Figure 1D

cancer_rules <-apriori(data, parameter = list(support =

0.01, confidence = 0.1, minlen = 2, maxlen =2))

KRAS_rules <-subset(cancer_rules, subset = lhs %ain% c(“KRAS”)) saveAsGraph((KRAS_rules), file=“KRAS_lhs_2-member_rules.graphml”)

\## Import and manipulate the saved file in Gephi

\# Figure 3A

cancer_rules <- apriori(data, parameter = list(support =

0.01, confidence = 0.1, minlen = 3))

KRAS_rules <- subset(cancer_rules, subset = lhs %ain% c(“KRAS”) &; count >= 17)

saveAsGraph((KRAS_rules), file=“KRAS_lhs_3-member_rules_count_17.graphml”)

\## Import and manipulate the saved file in Gephi

\# Figure 3B

cancer_rules <- apriori(data, parameter = list(support =

0.01, confidence = 0.1, minlen = 3))

KRAS_rules <- subset(cancer_rules, subset = lhs %ain% c(“KRAS”) & count >= 17)

\## Count >= 17, to include only rules that appear in at least 10% of KRAS-mutant samples

KRAS_PTPRs_rules <- subset(KRAS_rules, subset = rhs %ain% c(“PTPRs”))

inspect(KRAS_PTPRs_rules) ## Or write to a csv file

\# Figure 3C

cancer_rules <- apriori(data, parameter = list(support =

0. 01, confidence = 0.1, minlen = 3))

KRAS_rules <- subset(cancer_rules, subset = lhs %ain% c("KRAS") & count >= 17)

KRAS_STK11_rules <- subset(KRAS_rules, subset = rhs %ain% c("STK11"))

inspect(KRAS_STK11_rules) ## Or write to a csv file

